# Balance performance in aged mice is dependent on unipolar brush cells

**DOI:** 10.1101/2024.10.10.617602

**Authors:** Gabrielle Kizeev, Isabelle Witteveen, Timothy Balmer

## Abstract

The vestibular processing regions of the cerebellum integrate vestibular information with other sensory modalities and motor signals to regulate balance, gaze stability, and spatial orientation. A class of excitatory glutamatergic interneurons known as unipolar brush cells (UBCs) are highly concentrated within the granule cell layer of these regions. UBCs receive vestibular signals directly from primary vestibular afferents and indirectly from mossy fibers. Each UBC excites numerous granule cells and could contribute to computations necessary for balance-related motor function. Prior research has implicated UBCs in motor function, but their influence on balance performance remains unclear, especially in aged mice that have age-related impairment. Here we tested whether UBCs contribute to motor coordination and balance by disrupting their activity with chemogenetics in aged and young mice. Age-related balance deficits were apparent in mice > 6 months old. Disrupting the activity of a subpopulation of UBCs caused aged mice to fall off a balance beam more frequently and altered swimming behaviors that are sensitive to vestibular dysfunction. These effects were not seen in young (7-week-old) mice. Thus, disrupting the activity of UBCs impairs mice with age-related balance issues and suggest that UBCs are essential for balance and vestibular function in aged mice.

## Introduction

Accidental falls are the leading cause of injury and injury death among Americans >64 years old (Kakara et al., 2023). A major cause of accidental falls is an age-related decline in balance due to impaired sensorimotor function. Impaired balance in adults is typically a consequence of reduced vestibular function, somatosensation, vision, cognition, and muscle strength (Wagner et al., 2021). However, approximately half of the age-related balance impairment has been attributed to vestibular deficits in humans (Beylergil et al., 2019). Thus, understanding central mechanisms of sensory-motor function and vestibular processing are essential to develop strategies to overcome age-related balance impairments that can lead to injury, death, and disability.

The vestibular cerebellum, which includes lobules IX-X, the flocculus, and the paraflocculus, is a major site of vestibular processing. This region of the cerebellum integrates vestibular, visual, and other sensory modalities to coordinate movements, control balance, stabilize gaze, and maintain spatial orientation (Tarnutzer et al., 2011; Barmack and Pettorossi, 2021; Cullen, 2023). The cerebellar cortex receives sensory signals from primary vestibular afferents and secondary mossy fibers (Barmack, 2003). These signals excite granule cells, which modulate firing of Purkinje cells, the sole output of the cerebellar cortex. Purkinje cells inhibit the cerebellar nuclei, and those in the vestibular lobes also directly inhibit the vestibular nuclei that modulate lower motor neurons (Barmack, 2003; D’Angelo, 2018). In the vestibular cerebellum, unipolar brush cells (UBCs) are present in high densities, but their functional significance to vestibular processing has not been determined (Dino et al., 1999; Mugnaini et al., 2011). UBCs are excitatory interneurons that, like granule cells, receive vestibular signals directly from primary vestibular afferents and indirectly from mossy fibers (Dino et al., 2001; Balmer and Trussell, 2019). UBCs represent an additional processing layer that transforms and amplifies these signals to numerous granule cells and other UBCs (Mugnaini et al., 2011; Zampini et al., 2016; Balmer and Trussell, 2019; Hariani et al., 2023).

Disruption of Purkinje cell or cerebellar nuclei activity affects balance and motor performance, and is implicated in some genetically inherited forms of ataxia (Walter et al., 2006; Watase et al., 2008; Hourez et al., 2011; Todorov et al., 2012; Jayabal et al., 2016; Leto et al., 2016). The role of the granule cell layer in motor performance is less clear. In previous studies, approaches that disrupted large populations of granule cells had surprisingly little effect on balance performance. For example, increasing the activity of granule cells had no effect on motor performance or motor learning, measured using Rotor-rod, gait analysis, and eye blink conditioning, although non-motor effects were observed (Rudolph et al., 2020). To disrupt motor performance it was necessary to block neurotransmitter release by expressing tetanus toxin in granule cells (Yamamoto et al., 2003) or by deleting all three Ca_V_2 voltage-gated calcium channel genes simultaneously from granule cells (Galliano et al., 2013; Lee et al., 2023).

In contrast to granule cells, disrupting UBCs using various approaches has been reported to impair balance. In a TRPC3 gain of function mutant (Moonwalker), an early loss of UBCs correlates with the onset of ataxia and cannot be explained by the same mutation restricted to Purkinje cells alone (Becker et al., 2009; Sekerkova et al., 2013; Wu et al., 2019). In the same mutant mouse, compensatory eye movements were impaired, which may be the result of this loss of UBCs (Koops et al., 2023). Global knockout of an acid-sensing ion channel (ASIC5) that is present in a subpopulation of UBCs affected UBC excitability and caused motor impairment on Rotor-rod and balance beam tests (Kreko-Pierce et al., 2020). Finally, a study that ablated Golgi cells resulted in ataxia (Watanabe et al., 1998), but this approach presumably ablated UBCs as well, because it targeted all mGluR2-expressing cells, which includes UBCs (Neki et al., 1996; Jaarsma et al., 1998), so it remains unclear whether the ataxia was due to ablation of Golgi cells, UBCs, or both. The role of UBCs in animals challenged with age-related balance deficits has not been studied. The neural mechanisms that maintain balance performance in aging are especially important because falls are more common and more dangerous in the elderly.

In this study we disrupted the function of a subpopulation of UBCs, without affecting other types of cerebellar neurons, to test the hypothesis that these cells contribute to normal motor performance. We found that in aged mice (> 6 months old) chemogenetic manipulation of UBC activity impaired performance on a balance beam and altered swimming behaviors sensitive to vestibular function. These effects were not seen in young (7-week-old) mice. These data suggest that these cerebellar interneurons may maintain balance performance by compensating for age-related loss of vestibular function.

## Results

To determine the extent of age-related balance deficits in mice and whether UBCs contribute to balance behaviors, we utilized three behavioral tests: an accelerating Rotor-rod, a challenging balance beam, and a swimming test, in two age groups, young mice (7 weeks old) and old mice (>6 months old). We predicted that age-related vestibular deficits would be present in the old mice, because at this age C57BL/6 mice have evidence of hair cell loss in the vestibular periphery (Shiga et al., 2005). Habituation and training were required for the Rotor-rod and balance beam tests. After an initial habituation day that oriented the mice to the apparatuses (see Methods), there were 3 days of training in which mice performed three trials each day (Fig 1A). On the fourth day CNO was administered (Post-CNO) and the average performance across three trials was compared with that of the previous day (Pre-CNO). The results of these chemogenetic experiments are shown in Figures 3-4. The swimming test was performed once after CNO administration. Some animals performed all the behavioral tests, and these mice performed the swimming test last.

**Figure 1.**
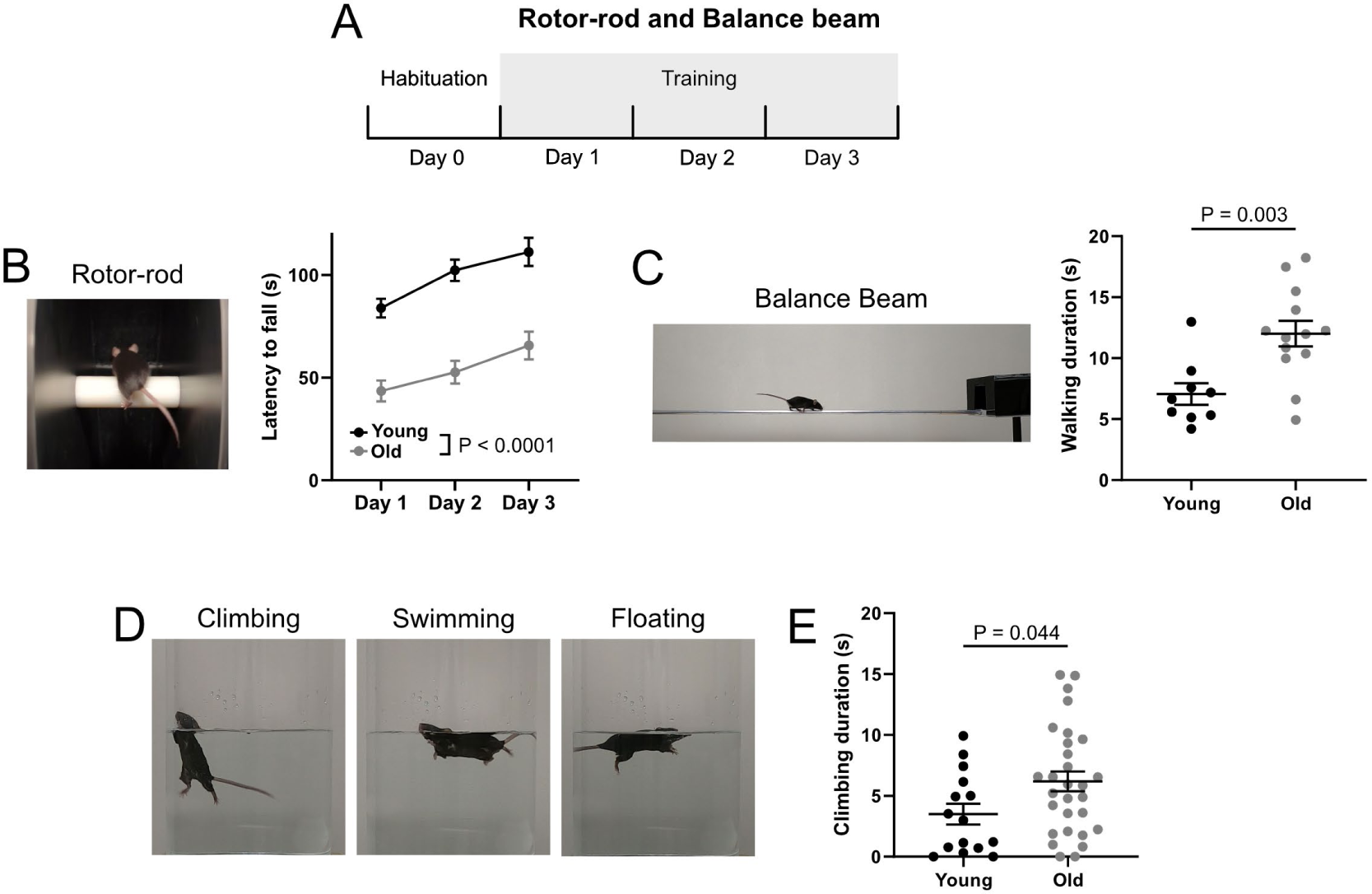
Age-related impairment in balance performance across three behavioral tests. A) Timeline showing the habituation and training schedule. Mice were habituated to the Rotor-rod and balance beam on Day 0 (see Methods). Training consisted of three trials per day for 3 days. To test whether old mice performed differently than young mice, performance during the 3-day training period was compared between age groups (Rotor-rod) or Day 3 performance was compared between groups (balance beam). B) An accelerating Rotor-rod was used to test the balance performance of mice in two age groups, young (7 weeks old) and old (> 6 months old). The average latency to fall was measured in three trials per mouse per day. Both groups were able to improve on the task over the three training days. Old mice fell off the rotating rod with a significantly shorter latency than did young mice. C) Mice were trained to cross a metal balance beam to reach a goal box. The walking duration, a measure of the latency to cross minus the time spent not walking, was significantly longer in the old mice. D) A swimming test was used to measure how the mice maintain their orientation in water. Examples of body positions during climbing, swimming and floating. E) Old mice climbed significantly more than young mice. Climbing is a measure reported to reflect vestibular dysfunction. These mice are wild-type littermate controls that received CNO injections.

### Age-related balance deficits in C57BL/6 mice

The Rotor-rod is a widely used balance test in which longer latencies to fall indicate better motor performance. Mice were able to balance on the accelerating rod for longer durations over the 3 training days in both age groups as a result of motor learning (Young: One-way RM ANOVA, F(1.91, 42.09) = 12.67, P < 0.0001, n = 23; Old: One-way RM ANOVA, F(1.82, 41.94) = 13.72, P < 0.0001, n=24, Fig 1B). The young mice had superior performance compared to the old group, as they were able to balance on the accelerating Rotor-rod longer (Young vs Old: Two-way RM ANOVA, F (1, 45) = 41.24, P< 0.0001, Fig 1B). This result supports the notion that balance is impaired in 6-month-old mice.

Motor performance was also tested on a balance beam consisting of an elevated ¼” diameter aluminum rod (Fig 1C). The animals were placed on the beam and were trained to walk across to a goal box that was dark and contained bedding material from their home cage. Training consisted of one habituation day followed by three training days during which each mouse performed the task three times (Fig 1A). Compared to the young mice, the old mice clearly had more difficulty crossing the beam, often moving along it with their bodies pressed to the beam rather than walking above it. When falls occurred the mouse was returned to the bar to continue to the goal box. The frequency of falls and walking duration (defined as the time spent walking along the beam not including pauses and falls, which is-a measure of moving speed), were used to quantify balance performance. The old mice moved along the beam more slowly than the young mice, resulting in a longer walking duration (Young: 7.06 ± 0.88 s (mean ± SEM), n = 9; Old: 12.00 ± 1.05 s, (mean ± SEM), n = 13; t-test, P = 0.003, Fig 1C). Although the old mice did not fall more frequently than the young mice (Young: 0.19 ± 0.08 falls per trial (mean ± SEM), n = 9; Old: 0.26 ± 0.10 falls per trial, (mean ± SEM), n = 13; t-test, P = 0.614), perhaps because of their careful and slow movements, it was clear that the aged mice had more difficulty walking along the balance beam than did the young mice.

A swimming test is a sensitive measure of vestibular dysfunction, presumably because in water somatosensory and proprioceptive cues are less informative and animals rely to a greater extent on their vestibular sense to stay upright (Ji et al., 2022). Thus, in a final behavioral test, we measured swimming behavior in young and old mice. Mice were placed in a cylinder of water and their behavior was video recorded and analyzed. We quantified the amount of climbing, swimming, and floating that the mice used to maintain their body position during the first minute in water. Climbing was defined as rapid paw movements that broke the surface of the water with the body oriented > 45° relative to the surface of the water; swimming was defined as rapid paw movements that did not break the surface with the body oriented < 45°; and floating was defined as slow or absent paw movements with a horizontal body position (Yuman et al., 2008). Examples of each behavior are shown in Figure 1D. In particular, increased climbing has been associated with vestibular dysfunction (Ji et al., 2022). We found that the old mice climbed for a significantly longer duration than did the young mice (Young: 3.50 ± 0.86 (mean ± SEM), n = 15; Old: 6.18 ± .082, (mean ± SEM), n = 29; unpaired t-test, P = 0.044, Fig 1C). Altogether, these three behavioral tests support the conclusion that mice over 6 months old exhibit impaired balance.

### CNO depolarizes GqDREADD-expressing UBCs in vitro

To test the role of UBCs in these balance behaviors we took a chemogenetic approach. A mouse line that expresses Cre recombinase (Cre) in UBCs (GRP-Cre) (Gerfen et al., 2013) was crossed to a reporter line that expresses hM3Dq designer receptor exclusively activated by designer drugs (GqDREADD) in the presence of Cre (Zhu et al., 2016), resulting in a line we refer to as GRP-DREADD, in which an IP injection of CNO increases the activity of a subpopulation of UBCs that accounts for ∼20% of ON UBCs (also referred to as Type II, or mGluR1α+) (Kim et al., 2012; Hariani et al., 2023). To confirm that genetically expressed GqDREADD affected the electrical activity of UBCs, GRP-DREADD mice were crossed with a Cre-dependent tdTomato reporter line (Ai9) (Madisen et al., 2010), resulting in offspring with expression of both GqDREADD and tdTomato in a subpopulation of UBCs that could be targeted for whole-cell recordings in acute brain slices (Fig 2A). Bath application of 10 µM CNO increased the spontaneous firing rate of these cells (Fig 2B-C), indicating that the CNO had an excitatory effect. In voltage-clamp, the application of CNO induced a large inward current (Fig 2D). In all the cells recorded, CNO induced an inward current that averaged -34.15 ± 18.75 pA (mean ± SD) (Fig 2E), a large depolarizing current for these small cells. These results demonstrate that this chemogenetic approach produced a reliable and disruptive effect on the GqDREADD-expressing UBCs.

**Figure 2.**
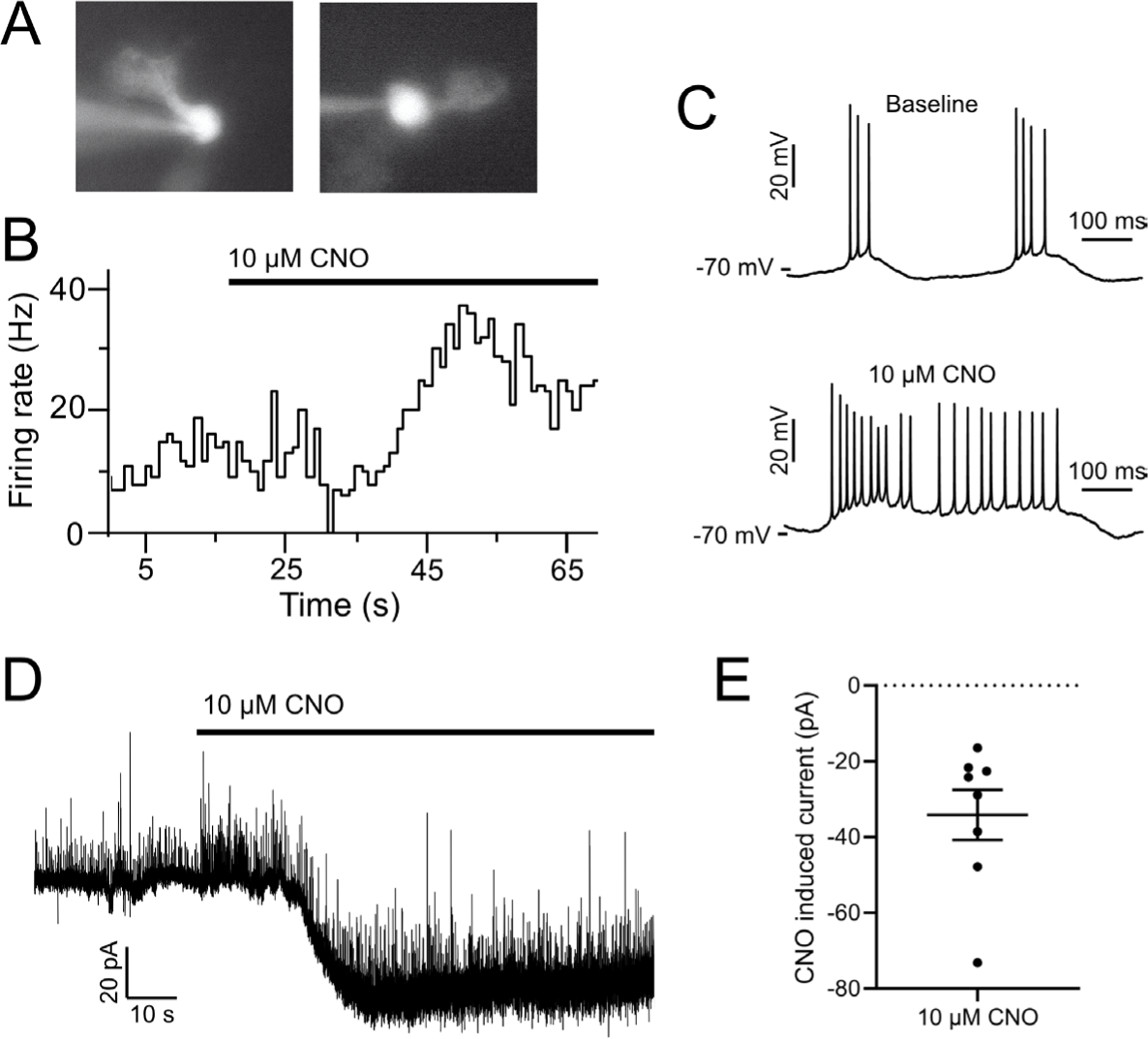
GqDREADD produces depolarizing current in GRP UBCs. A) Whole-cell recordings were made from genetically labeled UBCs in acute brain slices from GRP-Cre/GqDREADD/Ai9 mice. Two examples are shown. The recording pipette can be seen entering from the left. Dye fills confirmed UBC morphology. B) Bath application of CNO caused an increase in firing rate in current clamp. C) Example spontaneous spikes from before (top) and during bath application of CNO (bottom) in the experiment shown in B. D) In voltage clamp, CNO evoked a large inward current in an example UBC. E) In all UBCs tested, an inward current was produced by CNO. These are large depolarizing currents for cells of this size.

### Disruption of UBC activity did not impair aged mice on Rotor-rod

GRP-DREADD mice were indistinguishable from their littermate controls in their general posture and locomotor behaviors. After an initial habituation day, each mouse was tested 3 times per day for 3 days, followed by a fourth day during which all animals were injected with Clozapine-N-oxide (CNO-3 mg/kg) > 30 minutes prior to testing (Fig 3A). Average latencies to fall over the three trials were compared with those of the previous day. Control mice were littermates that were negative for the Cre or GqDREADD genes and received the same CNO injection to control for potential off-target effects of CNO (Manvich et al., 2018). Although there was a main effect of age on latency to fall (3-way RM ANOVA, F(1,89) = 38.49, P < 0.0001), as seen in Figure 1B, no differences were observed between GRP-DREADD mice or their littermate controls within age groups (3-way RM ANOVA, no significant main effect of CNO, genotype, or interactions Fig 3B-D) indicating that disrupting UBC activity did not affect their ability to balance on the Rotor-rod in mice of either age.

**Figure 3.**
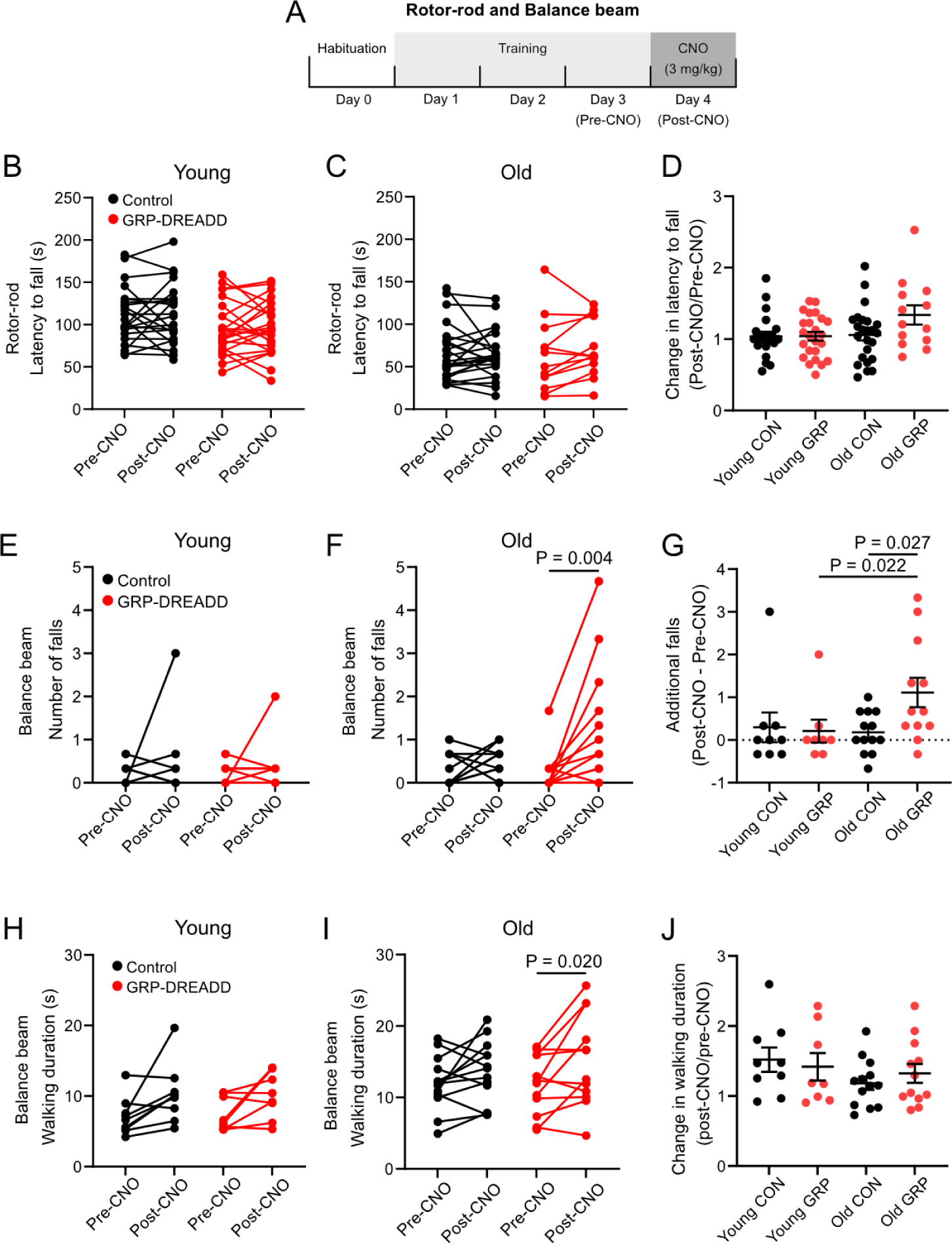
Effect of disruption of UBCs on balance performance in young and aged mice. A) Timeline showing the habituation, training, and testing schedule. Mice were habituated to the Rotor-rod and balance beam on Day 0 (see Methods). Training consisted of three trials per day for 3 days. To test whether old mice performed differently than young mice, performance during the 3-day training period was compared between age groups (Rotor-rod) or Day 3 performance was compared between groups (balance beam). On Day 4, CNO was injected IP (3 mg/kg), and performance was compared within subjects between Day 3 (Pre-CNO) and Day 4 (Post-CNO). B) Mice were trained on accelerating Rotor-rod for 3 days. Their performance was compared between their third training day (Pre-CNO) and the following day after CNO treatment (Post-CNO). C) Rotor-rod performance of old mice on the third training day (Pre-CNO) and the following day after CNO treatment (Post-CNO). D) Comparison of the effect of CNO on latency to fall off the Rota-rod across groups. E) Mice were trained to cross a metal balance beam to reach a goal box. The number of falls were averaged across three trials on the third day of training (Pre-CNO) and compared to the average number of falls during three trials on the following day after CNO treatment (Post-CNO). CNO treatment did not affect the number of falls in young control. F) CNO treatment did not affect the number of falls in the old control group, but significantly increased the number of falls in the old GRP-DREADD group. G) Mean number of additional falls after CNO treatment. The old GRP-DREADD mice fell more after CNO compared to before CNO and this was a larger effect than the effect seen in old littermate controls and young GRP-DREADD mice. H) Walking duration was not affected by CNO in young mice of either genotype. I) The walking duration of old GRP-DREADD mice was longer after CNO treatment compared to before. J) Comparison of the effect of CNO on walking duration across groups.

### Disruption of UBC activity impaired the performance of aged mice on a balance beam

The Rotor-rod has been reported to be less sensitive to vestibular deficits than other behavioral tests including balance beam and swimming tests (Stanley et al., 2005; Luong et al., 2011; Ji et al., 2022). Thus, motor performance was tested on a balance beam consisting of an elevated ¼” diameter aluminum rod. Walking duration was measured as described above. In many cases, the mice slipped around the beam to an upside-down position or fell off entirely, both of which were considered falls. When falls occurred the mouse was returned to the bar to continue to the goal box. The frequency of these falls provided a robust and easily scored measure of balance performance. On the fourth day, >30 min after CNO injections, the mice were tested 3 times and their average number of falls per trial was compared to those of the previous day.

In the young group, chemogenetic disruption of UBC activity had no significant effect on the number of falls of the GRP-DREADD mice or the littermate controls (Young Control: Pre-CNO vs Post-CNO, Wilcoxon matched-pairs signed rank test, P = 0.8125, n = 9; Young GRP-DREADD: Pre-CNO vs Post-CNO, Wilcoxon matched-pairs signed rank test, P = 0.999, n = 8, Fig 3E). In contrast, in the old group CNO treatment increased the number of falls in the GRP-DREADD mice compared to the previous day (Old GRP-DREADD Pre-CNO vs Post-CNO, Wilcoxon matched-pairs signed rank test, P = 0.004, n = 12), but had no significant effect on the littermate controls (Old Control Pre-CNO vs Post-CNO, Wilcoxon matched-pairs signed rank test, P = 0.266, n = 13, Fig 3F). The effect of CNO was significantly greater in the old GRP-DREADD group than the old Control group (effect of CNO on old GRP-DREADD: 1.11 ± 0.34 more falls (mean ± SEM), n = 12; effect of CNO on old Controls: 0.18 ± 0.13 more falls (mean ± SEM), n = 13; Mann-Whitney test, P = 0.027), as well as being significantly greater than the young GRP-DREADD group (Mann-Whitney test, P = 0.022) (Fig 3G).

The walking duration was compared using a 3-way RM ANOVA, which showed significant main effects of age (P=0.0005) and CNO (P = 0.0001), but no significant main effects of genotype or interactions across factors. Planned within-subject comparisons indicated that the walking duration of the old GRP-DREADD mice was significantly longer after CNO treatment (Old GRP-DREADD Pre-CNO: 11.82 ± 4.15 s (mean ± SEM), Old GRP-DREADD Post-CNO: 15.25 ± 6.44 s (mean ± SEM), n = 12; Sidak’s multiple comparisons test, P = 0.020, Fig 3I). However, the size of the effect of CNO on old GRP-DREADD mice was not significantly greater than the effect of CNO on old littermates (Proportional effect of CNO on old GRP-DREADD: 1.324 ± 0.1360 s (mean ± SEM), n = 12; Proportional effect of CNO on old Control: 1.182 ± 0.0943 s (mean ± SEM), n = 13; unpaired t-test, P = 0.394, Fig 3J). Taken together, the balance beam tests revealed that disrupting UBC function impairs balance in aged mice, but not young mice.

### Disruption of UBC activity alters swimming behaviors in aged mice

Swimming behavior can reveal vestibular dysfunction that is not detected in other behavioral tests. Mice were placed in a cylinder of water and their swimming behavior was video recorded and analyzed. We quantified the durations of climbing, swimming, and floating that the mice used to maintain their body position during the first minute in water. Examples of each behavior are shown in Figure 1D. In the young mice, no significant differences in the duration nor the number of occurrences of any of these behaviors were seen between the experimental and control groups after CNO treatment (Fig 4A-B). In contrast, disruption of UBC activity in the old mice caused them to climb for a shorter total duration than did the CNO-injected littermate controls (Mann-Whitney test, P = 0.0002, Fig 4C). In addition, the CNO-treated GRP-DREADD mice made fewer climbing attempts than did the controls (GRP-DREADD: 5.66 ± 0.70 climbing bouts, n = 30; Control: 2.83 ± 0.48 climbing bouts, n = 29; unpaired t-test, P = 0.003) and also swam less frequently (GRP-DREADD: 6.13 ± 0.45 swimming bouts, n = 30; Control: 8.21 ± 0.71 swimming bouts, n = 29; unpaired t-test, P = 0.016) and instead floated more frequently (GRP-DREADD: 2.97 ± 0.35 floating bouts, n = 30; Control: 1.93 ± 0.26 floating bouts, n = 29; Mann-Whitney test, P = 0.036) (Fig 4D). The old CNO-treated GRP-DREADD mice appeared uncoordinated in their swimming. In particular, they more often rolled sideways enough to necessitate paddling to recover to a stable position, and after bumping into the wall they frequently paused and appeared disoriented. Overall, these results suggest that swimming behaviors of aged mice are affected by the chemogenetic disruption of UBCs and implicate these cerebellar interneurons in vestibular function.

**Figure 4.**
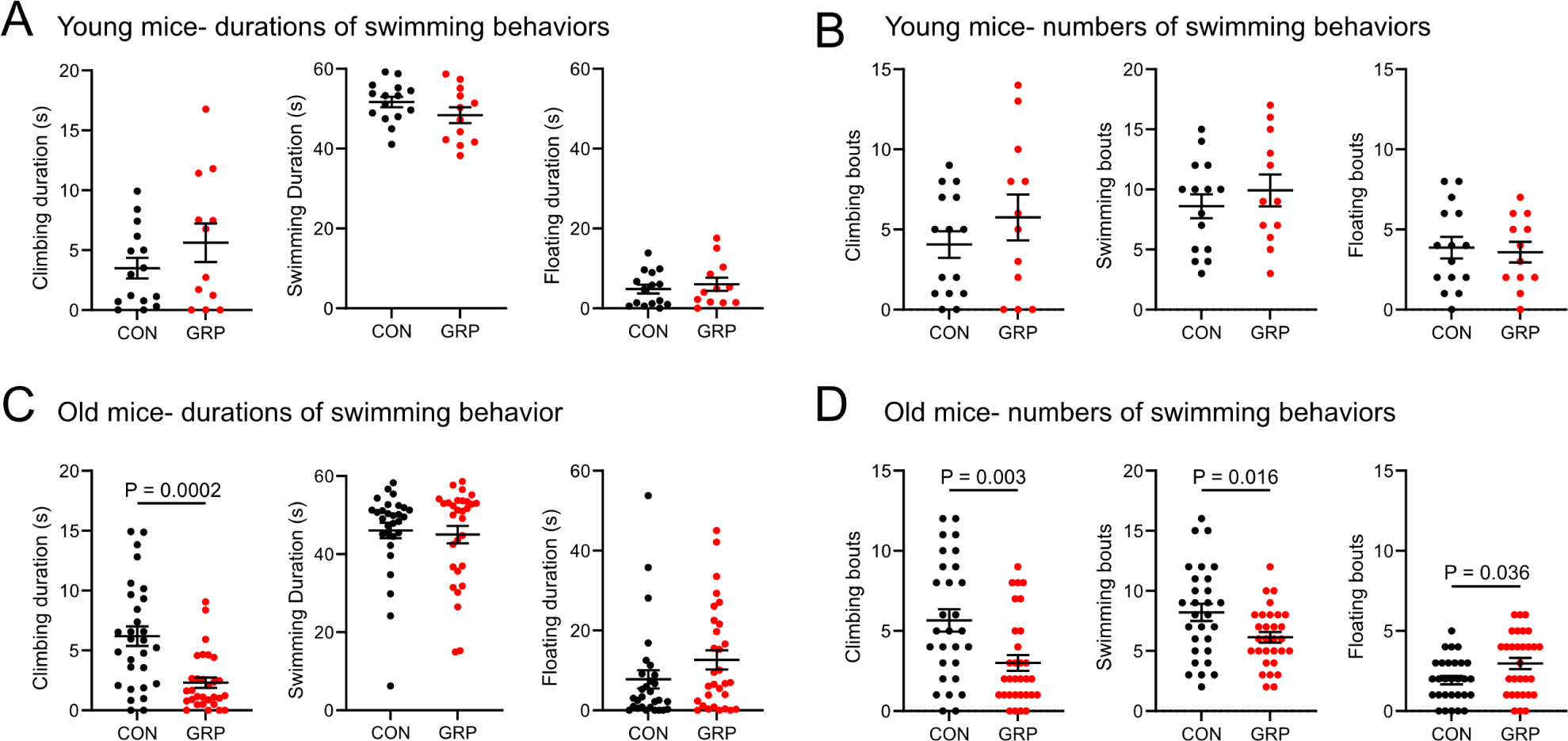
Disruption of UBC function affects swimming behaviors in old mice. A) A swimming test was used to measure how the mice maintain their orientation in water during the first minute. All mice received CNO injections. In the young mice, there were no significant differences in the duration of climbing, swimming, or floating between GRP-DREADD mice and littermate controls. B) In the young mice, there were no significant differences between groups in the number of occurrences of these behaviors during the test. C) Durations of swimming behaviors of old mice during the first minute of the test. In contrast to the young mice, the old GRP-DREADD mice climbed significantly less than their littermate controls. D) Old GRP-DREADD mice performed fewer bouts of climbing and swimming, while floating more frequently, than did the littermate controls.

## Discussion

The vestibulocerebellum is involved in balance, posture, compensatory eye movements, and the generation of an internal model of self-motion (Cullen, 2023). Despite their prevalence in vestibular cerebellar regions, the role of UBCs in cerebellar function remains unclear. Here we disrupted the activity of a subpopulation of UBCs and measured the effect on balance behaviors across different ages. We found that chemogenetic disruption of UBCs caused old mice (> 6 months) to fall off a balance beam more frequently and affected their swimming behavior. In the young (7 weeks) age group, the same chemogenetic disruption of UBC activity did not produce significant effects in either test. We confirmed that the chemogenetic manipulation disrupted UBC activity using acute brain slice electrophysiology. Based on these results we propose a model in which UBCs may maintain vestibular function by amplifying residual signals that are reduced in aged mice.

### UBCs may compensate for degraded peripheral input in aged mice

Chemogenetic disruption of UBCs affected motor performance in aged mice but not young mice. One potential explanation for this result is that young brains may be more flexible and can compensate for the disruption of a small population of neurons and that aged brains cannot. There is some evidence that there are mechanisms that prevent age-related sensory deficits from disrupting vestibular function. For example, despite evidence for loss of hair cells in the semicircular canals of 6-month-old C57BL/6 mice, there are no apparent effects on vestibular ocular reflex function, which contrasts with age-related loss of cochlear hair cells that coincides with hearing loss in the same animals (Shiga et al., 2005). ON UBCs receive direct input from semicircular canal afferents and could enhance vestibular signals that may be diminished due to age-related hair cell loss or peripheral damage (Balmer and Trussell, 2019). One synaptic mechanism that may be utilized by ON UBCs to enhance signals is their unusual AMPA receptor mediated response to glutamate that extends the duration of their synaptic input (Rossi et al., 1995; Lu et al., 2017; Balmer et al., 2021). UBCs provide this extended duration signal to numerous downstream targets via their branching axons to hundreds of granule cells, as well as other UBCs that do the same (Nunzi and Mugnaini, 2000; van Dorp and De Zeeuw, 2015; Hariani et al., 2023). Thus, UBCs could theoretically compensate for peripheral loss of function by enhancing the remaining input. Synaptic plasticity occurs at numerous points in the circuit and may compensate for age-related changes as well.

Circuit mechanisms including inhibition of parts of network that no longer receive accurate signals may also contribute to the maintenance of normal function in aged animals. For example, Golgi cells affect the temporal dynamics of the cerebellar cortical circuit and modulate the activity of granule cell ensembles to improve pattern separation (Rossi et al., 2003; Kanichay and Silver, 2008; Fleming et al., 2024). Granule cell ensembles that no longer provide accurate vestibular information could be specifically inhibited. In aged mice these mechanisms may be especially important for normal function and are thus more sensitive to chemogenetic disruption than are young mice. Further experiments are warranted to test how synaptic and circuit mechanisms prevent or delay age-related vestibular and balance impairment.

### The role of different subtypes of UBCs

UBCs can be classified by their differential protein expression (Nunzi et al., 2002, 2003; Mugnaini et al., 2011; Kim et al., 2012) or by their electrophysiological response to glutamate (Borges-Merjane and Trussell, 2015). Single-cell RNA sequencing studies have revealed that, despite the apparent difference in protein expression, RNA expression for the same genes varies continuously across the population of UBCs (Guo et al., 2021; Kozareva et al., 2021). Nonetheless, mouse lines have been generated that genetically label subpopulations of UBCs with distinct properties (Kim et al., 2012; Hariani et al., 2023). The GRP-Cre line used here expresses Cre in ∼20% of the ON UBC population that is excited by glutamate due to AMPA and mGluR1 mediated currents (Hariani et al., 2023). The result reported here that only old mice were impaired by GRP UBC disruption may be related to the low proportion of UBCs affected. Indeed, in a study that disrupted the electrical activity of a larger population of ON UBCs, balance was disrupted in young mice (Kreko-Pierce et al., 2020). It is possible that there is more redundancy or parallel processing in young mice that can be used to compensate for disordered activity in part of the circuit, which is not available in old mice. Disrupting the activity of OFF UBCs that are inhibited by glutamate may also be informative. UBC subtypes may affect the temporal processing of vestibular input in different ways and comparing the behavioral or neurophysiological results caused by subtype-specific manipulations may improve our understanding of their distinct roles.

### Behavioral tests of vestibular and balance function in mice

Measuring vestibular function in mice is challenging. In some cases, mutations that profoundly alter the vestibular periphery can have subtle effects in behavioral tests (Shiga et al., 2005; Ji et al., 2022). Perhaps this is not surprising given the remarkable compensation that occurs in the central nervous system to prevent impairment, for example, during chronic vestibulocerebellar disease or during recovery from unilateral labyrinthectomy (Smith and Curthoys, 1989; Tarnutzer et al., 2011). To measure what was expected to be a small vestibular effect given the small proportion of UBCs affected by our chemogenetic manipulation, we utilized the Rotor-rod, a commonly used assessment of gross motor performance and coordination, as well as a balance beam and a swimming test that has detected vestibular impairment in previous work (Tung et al., 2016). To further enhance the sensitivity of the balance beam test, we used a smooth aluminum beam, which was more slippery than the more commonly used wooden beam. Mice appeared to find it challenging and fell routinely, either spinning to an upside-down position or falling off entirely. This allowed an unequivocal measure of balance that we favor over foot slips that are a common measure on a wooden balance beam test. Swimming behaviors have become a more common measure for vestibular dysfunction in mice. One advantage of the swimming test is that no training is required. A disadvantage is that a within subject design (before vs after a manipulation) is not possible, because the task is novel only on the first test and a second test would likely be different after the mouse learns to float.

Mice over 6 months old had more difficulty on the Rotor-rod than young mice, measured as shorter latencies to fall, which agrees with previous Rotor-rod experiments in aged mice (Serradj and Jamon, 2007; Tung et al., 2016). However, there were no effects of CNO on Rotor-rod performance in any age group, despite differences on balance beam performance and swimming behavior. Similar results were reported in another study in which mutants with altered utricle hair cell polarity and impaired bidirectional vestibular sensitivity performed normally on the Rotor-rod, whereas balance beam and swimming behaviors revealed deficits (Ji et al., 2022). In that study, the mice with peripheral vestibular defects climbed more than the controls and the authors describe a frantic swimming phenotype. CNO treatment may have impaired coordination or spatial orientation in a different way in the present study. Although the old mice climbed more than young mice, old CNO treated GRP-DREADD mice climbed less than did their CNO treated littermate controls. We observed that the old CNO treated GRP-DREADD mice appeared to spend more time paddling on one side to maintain their orientation and to avoid rolling sideways than did the littermate controls. Thus, the phenotype here may be due to an effect on the animals’ perception of their orientation in space that does not lead to frantic swimming. These results support the conclusion that swimming behaviors are sensitive to the disruption of UBC function.

### Chemogenetic approach disrupted UBC processing

We confirmed in vitro that CNO application depolarizes UBCs in our GRP/GqDREADD mouse line. These inward currents were tens of picoamperes, which for a cell with an average input resistance of over 500 MΩ (Hariani et al., 2023), produces a significant depolarization. The effect of depolarization is not likely to be a simple increase in firing rate in these cells. UBCs have been reported to fire in bursts or with a regular pattern, depending on their resting membrane potential (Diana et al., 2007). Chemogenetic depolarization of a few millivolts could therefore change the firing pattern of the cell profoundly. Depolarization could cause a phase shift in the temporal response of these cells, which could cause a timing mismatch of vestibular and other input in the vestibulocerebellar network that could lead to motor impairment.

CNO has been reported to be reverse metabolized into the psychoactive drug clozapine and may have other off-target effects (Manvich et al., 2018). We do not anticipate that at the concentration used here this is a significant concern, but nonetheless, we also injected CNO into the littermates that lacked the DREADD or Cre genes to control for these effects. About half of each cohort were controls and they were run together with the experimental animals on the same days. The experimenters were blind to the genotype of the animals both during the experiments and during analysis.

### Limitations of this study

The GRP-cre mouse line is specific for a subpopulation of UBCs and does not express Cre in other cerebellar neurons, although Cre expression is present in some other neurons including neurons in CA3 of the hippocampus and the superficial dorsal horn. Although we cannot completely exclude the possibility that these neurons could have contributed to the effects that were observed, cerebellar and not hippocampal lesions disrupt motor coordination (Goddyn et al., 2006) and specific ablation of the Cre expressing neurons in the dorsal horn of this mouse line had no effect on Rotor-rod performance in a previous study (Albisetti et al., 2019).

This study did not address the role of UBCs in motor learning, as the chemogenetic manipulation of activity was done after a training period in the Rotor-rod and balance beam experiments. The notion that granule cells are essential for motor learning is supported by numerous studies and theoretical models (Marr, 1969; Albus, 1971; Medina and Mauk, 2000; Ito, 2006; Dean et al., 2010) and the potential role of UBCs in orchestrating their activity to support motor learning is an important avenue for future research.

## Conclusion

Vestibular function is essential to an animal’s survival and, perhaps for this reason, is robust. However, age-related vestibular impairment compounds other sensorimotor deficits, which results in reduced balance performance. This can lead to falls which are a leading cause of injury and disability. The behavioral effects we observed after disrupting UBC processing could be due to impaired sensorimotor integration, inaccurate internal models of self-motion, or unsteady eye movements during head motion. Recent work has suggested a role for UBCs in the flocculus in compensatory eye movements (Koops et al., 2023). Although reflexive eye movement may be important to stabilize vision during head movements while an animal walks on a beam, the differences in swimming behaviors after UBC disruption is more consistent with an impaired sense of spatial orientation. Further experiments are necessary to differentiate between these explanations and would improve our understanding of the role of UBCs in vestibular processing.

## Methods

### Animals

Mice of both sexes in roughly equal numbers were used from the following mouse lines and their crosses: GRP-Cre: Tg(Grp-Cre)KH107Gsat (MMRRC_031182-UCD) (Gerfen et al., 2013), tdTomato reporter (Ai9): Gt(ROSA)^26Sortm9(CAG-tdTomato)Hze^ (IMSR_JAX:007909) (Madisen et al., 2010), GqDREADD reporter: Tg(CAG-CHRM3*,-mCitrine)1Ute (IMSR_JAX:026220) (Zhu et al., 2016). Mice were bred in a colony maintained in the animal facility managed by the Department of Animal Care and Technologies and all procedures were approved by Arizona State University’s Institutional Animal Care and Use Committee under protocol #21-1817R. Mice were provided food and water ad libitum and housed in Thoren ventilated caging, at a temperature of 23 ± 1°C and a 12 h light/12 h dark cycle. Behavioral testing was performed during the light cycle. Mice were genotyped by PCR.

### Acute brain slice electrophysiology

Mice were anesthetized with isoflurane and decapitated. The brain was rapidly extracted into an ice-cold high-sucrose artificial cerebral spinal fluid (ACSF) solution containing (in mM): 87 NaCl, 75 sucrose, 25 NaHCO3, 25 glucose, 2.5 KCl, 1.25 NaH2PO4, 0.5 CaCl2, and 7 MgCl2, bubbled with 5% CO2/95% O2. Parasagittal cerebellum sections containing the nodulus were cut at 300 µm thickness with a vibratome (7000smz-2, Campden Instruments) in ice-cold high-sucrose ACSF. Immediately after cutting, slices were incubated at 35°C recording ACSF for 30–40 minutes, followed by storage at room temperature. Recording ACSF contained (in mM): 130 NaCl, 2.1 KCl, 1.2 KH2PO4, 3 Na-HEPES, 10 glucose, 20 NaHCO3, 2 Na-pyruvate, 1.5 CaCl2, 1 MgSO4, and 0.4 Na-ascorbate. Solutions were bubbled with 5% CO2/95% O2 (300–305 mOsm).

Slices were transferred to a submerged recording chamber and perfused with ACSF heated to 33-35°C with an inline heater at 3 ml/min. Slices were viewed using an infrared Dodt contrast mask and a 60X water-immersion objective and an IR-2000 camera (Dage-MTI) on an Olympus BX51 fixed stage microscope. In the GRP-Cre/Ai9/GqDREADD slices, ON UBCs were identified by their tdTomato fluorescence. Pipettes were pulled from borosilicate 1.2 mm OD glass capillaries (A-M Systems) to a tip resistance of 5–8 MΩ. The internal pipette solution contained (in mM): 113 K-gluconate, 9 HEPES, 4.5 MgCl2, 0.1 EGTA, 14 Tris-phosphocreatine, 4 Na2-ATP, and 0.3 Tris-GFP, with osmolality adjusted to ∼290 mOsm with sucrose and pH adjusted to pH 7.3 with KOH. Reported voltages were corrected for a 10-mV liquid junction potential. Whole-cell recordings were amplified (10X), low-pass filtered (10 kHz Bessel, Multiclamp 700B, Molecular Devices), and digitized using pClamp 11 software (20-50 kHz, Digidata 1550B, Molecular Devices). Further digital filtering was performed offline. Cells were voltage-clamped at -70 mV. CNO (10 µM) was added to the bath to test the response of the UBCs expressing the GqDREADDs.

### Rotor-rod

The experimental procedure for the Rotor-rod (San Diego Instruments) was based on previous protocols (Carter et al., 2001; Deacon, 2013; Tung et al., 2014; Kreko-Pierce et al., 2020). Each day the mice were transferred to the experimental room in their home cage at least 10 minutes before starting the task. The first day was a habituation day that allowed the mice to become familiar with the apparatus, followed by three training days and a final test day. On the habituation day the mice were placed on the stationary rod (∼3 cm diameter) for a total of one minute, replacing them on the rod if they fell. The Rotor-rod was then set to a constant speed of 4 RPM and the mice were placed on the rod while it was stationary and, once facing the appropriate direction, the rotation was initiated and maintained for 300 s. If the mouse fell, it was placed back on the rod up to three times. For all 4 days after the habituation day an accelerating protocol was used, in which the rotation was 4 RPM for the first 10 s and then increased to 40 RPM linearly over the remaining 300 s. Mice were placed on the stationary rod and once they faced the appropriate direction the accelerating protocol was initiated. When the mice fell, the software automatically recorded the latency to fall in seconds. The mice were returned to their holding cage for an intertrial period of at least 5 minutes. Each mouse was given three trials per day over the three consecutive training days. If the mouse fell within the first 10 seconds (i.e., before the beginning of the acceleration), then the trial was re-done. Between each animal and at the end of the day’s session the apparatus was cleaned with 70% ethanol followed by Odormute. On the fourth day (test day), both experimental mice (GRP-GqDREADD) and control mice (littermates lacking Cre and/or GqDREADD genes) were given an IP injection of CNO (3 mg/kg, Tocris, Cat. No. 6329, 0.5 mg/ml in saline) >30 minutes before the assessment but was otherwise identical to the training days.

### Balance beam

The experimental procedure for the balance beam was based on previous protocols (Carter et al., 2001; Luong et al., 2011; Deacon, 2013; Kreko-Pierce et al., 2020). One modification to these protocols is that a smooth ¼” diameter aluminum balance beam was used, which we found to be superior to a less challenging wooden beam, because in contrast to foot slips that are a typical measure of motor errors on a wooden beam, falls are unambiguous and easily scored. The balance beam was supported by two stands and led to a goal box that was 16 x 10 x 9 cm. The beam was mounted ∼40 cm above a foam pad. Lines were drawn on the beam to demarcate an 80-cm span with an additional 10-cm starting region and a 10-cm ending region, providing both an on-ramp and an off-ramp for the mouse. These 10-cm spans on either side of the 80-cm length were designed to reduce the effect of cautious starts and entries into the goal box, which often occurred because the mouse investigated the apparatus at these points.

Each day the mice were transferred to the experimental room in their home cage at least 10 minutes before starting the task. A small piece of nesting material from the mouse’s home cage was placed in the goal box. If the mouse stopped walking along the beam, it was encouraged to continue with a gentle nudge. This was necessary for mice that had stalled on the beam, in many cases to groom or to sniff the air and occurred with sufficient frequency that we found that the total time to cross the beam that included these pauses was unrelated to balance performance. The apparatus was cleaned with 70% ethanol followed by Odormute between each mouse.

On the habituation day each mouse was given three consecutive trials on a ¼” diameter wooden beam, with an intertrial period of 5-10 seconds in the goal box. The three training days were identical, each having three trials for each mouse. As described for the Rotor-rod above, on the fourth day (test day), both experimental mice (GRP-Cre/ GqDREADD) and control mice (littermates lacking either the Cre or GqDREADD gene) were given an IP injection of CNO (3 mg/kg, Tocris, Cat. No. 6329, 0.5 mg/ml in saline) >30 minutes before the assessment but was otherwise identical to the previous 3 training days. A video camera (GoPro HERO8) was used to record each trial for post hoc analysis at 1080 X 1920 resolution and 60 frames per second. The videos were manually annotated with timestamps indicating the beginning and end of each behavior including walking, pausing, falling/replacement on the bar, by an experimenter that was blind to the genotype of the animal in Adobe Premiere. The same scorer (I.W.) annotated all the animals to maintain consistency. These annotations were exported as a text file and processed in Matlab (Mathworks).

### Swimming test

The swimming test was used to test vestibular function and was based on previous research that has shown that this is an sensitive behavioral assay (Gray et al., 1988; Balaban, 2002; Ji et al., 2022). Mice were transferred to the experimental room in their home cage at least 10 minutes before starting the task. The apparatus consisted of a transparent, 30 cm tall, 20 cm diameter acrylic cylinder filled with room temperature (24 ± 1°C) water to a level that prevented them from touching the bottom (∼20 cm deep). Experimental mice (GRP-DREADD) and control mice (littermates lacking either the Cre or GqDREADD gene) were given an IP injection of CNO (3 mg/kg, Tocris, Cat. No. 6329, 0.5 mg/ml in saline) >30 minutes before the assessment. Mice were held 5 cm over the water by their tail and released into the container. Their swimming behaviors were recorded from the side, parallel to the waterline, with a video camera (GoPro HERO8) for 6 minutes. Climbing was defined as rapid paw movements that broke the surface of the water with the body oriented > 45° relative to the surface of the water; swimming was defined as rapid paw movements that did not break the surface with the body oriented < 45°; and floating was defined as slow or absent paw movements with a horizontal body position (Yuman et al., 2008). After the 6-minute trial the mouse was retrieved from the water, patted dry with a paper towel, and returned to its cage. Each mouse was observed for 10 minutes to ensure it was active and grooming itself. The videos were manually annotated with timestamps indicating the beginning and end of each behavior including swimming, climbing, and floating by and experimenter that was blind to the genotype of the animal in Adobe Premiere. The annotations were exported as a text file and processed in Matlab (Mathworks) to calculate the duration and number of bouts of each behavior during the first minute. The same scorer (I.W.) annotated all the animals to maintain consistency.

### Statistical analysis

Data within groups were tested for normality using D’Agostino & Pearson tests. If the groups being compared were normally distributed, unpaired t-tests were used, otherwise Mann-Whitney tests were used. Within-subjects comparisons used one-, two-, or three-way repeated measures ANOVAs, followed by Sidak’s multiple comparisons tests, where data within all groups were normally distributed. Otherwise, Wilcoxon matched-pairs signed rank tests were used for within-subject comparisons. In cases where within-subject comparisons showed significant differences in one group and not another, a result that itself does not suggest a significant difference in the effect between groups (Gelman and Stern, 2006; Nieuwenhuis et al., 2011), a direct statistical comparison of the effect of the manipulation (e.g. CNO) between groups was made, using either unpaired t-tests or Mann-Whitney tests. Prism (GraphPad) software was used for statistical analyses.

## Acknowledgements

We thank Dr. Luis Martinez (Trinity College) for help with statistical analyses and members of the lab for help with behavioral analyses.

## Declarations

### Ethical Approval

All animal procedures were approved by Arizona State University’s Institutional Animal Care and Use Committee under protocol #21-1817R.

### Funding

Funding was provided by the NIH/NIDCD R00 DC016905, Hearing Health Foundation, and National Ataxia Foundation.

### Availability of data and materials

All data supporting the findings of this study are available within the paper.

## Notes

### Competing Interest Statement

The authors have declared no competing interest.

